## Supplementary figures and images for "The T Cell Receptor β Chain Repertoire of Tumor Infiltrating Lymphocytes Improves Neoantigen Prediction and Prioritization"

### Supplementary Figure 1

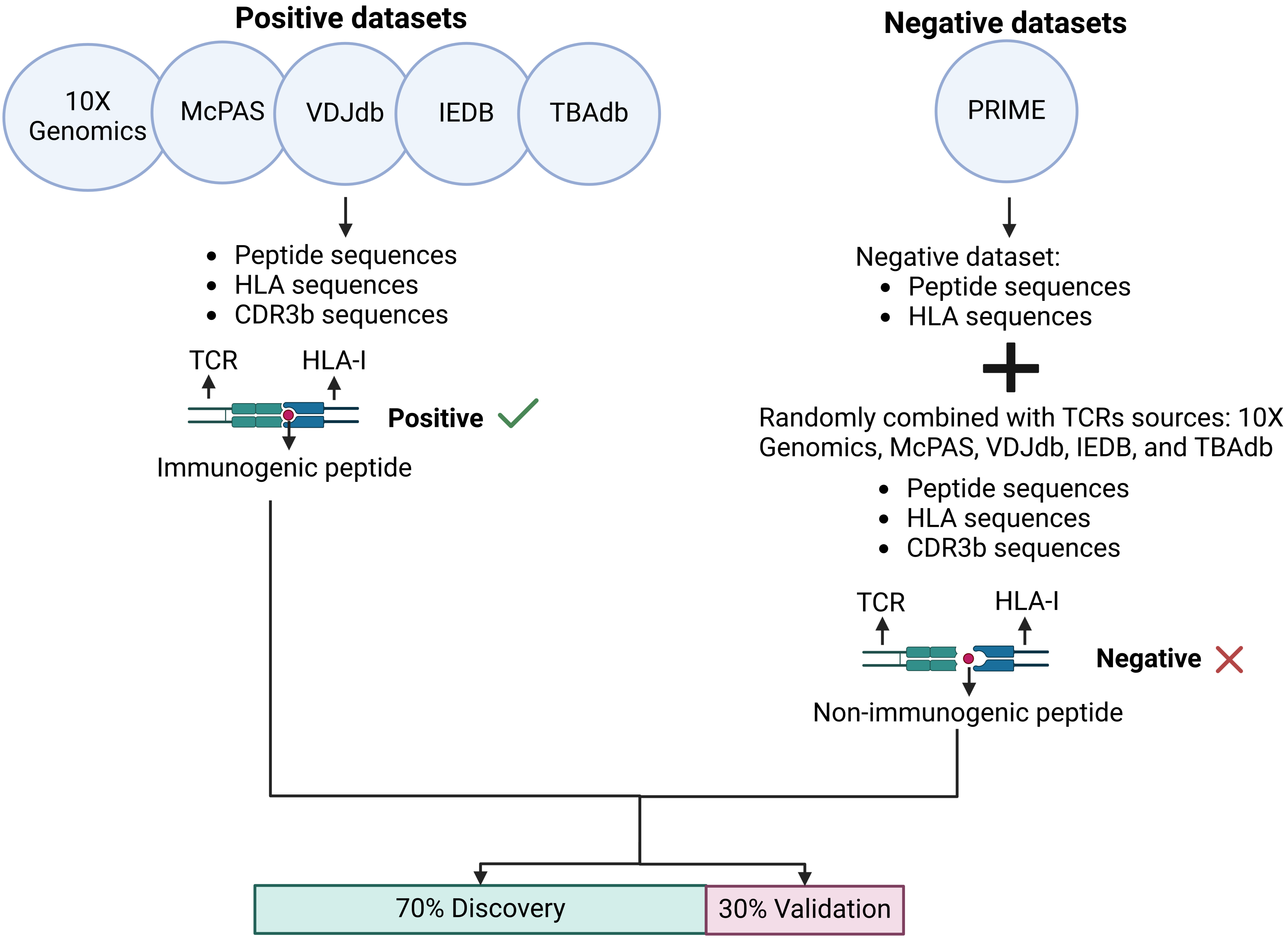

### Supplementary Figure 2

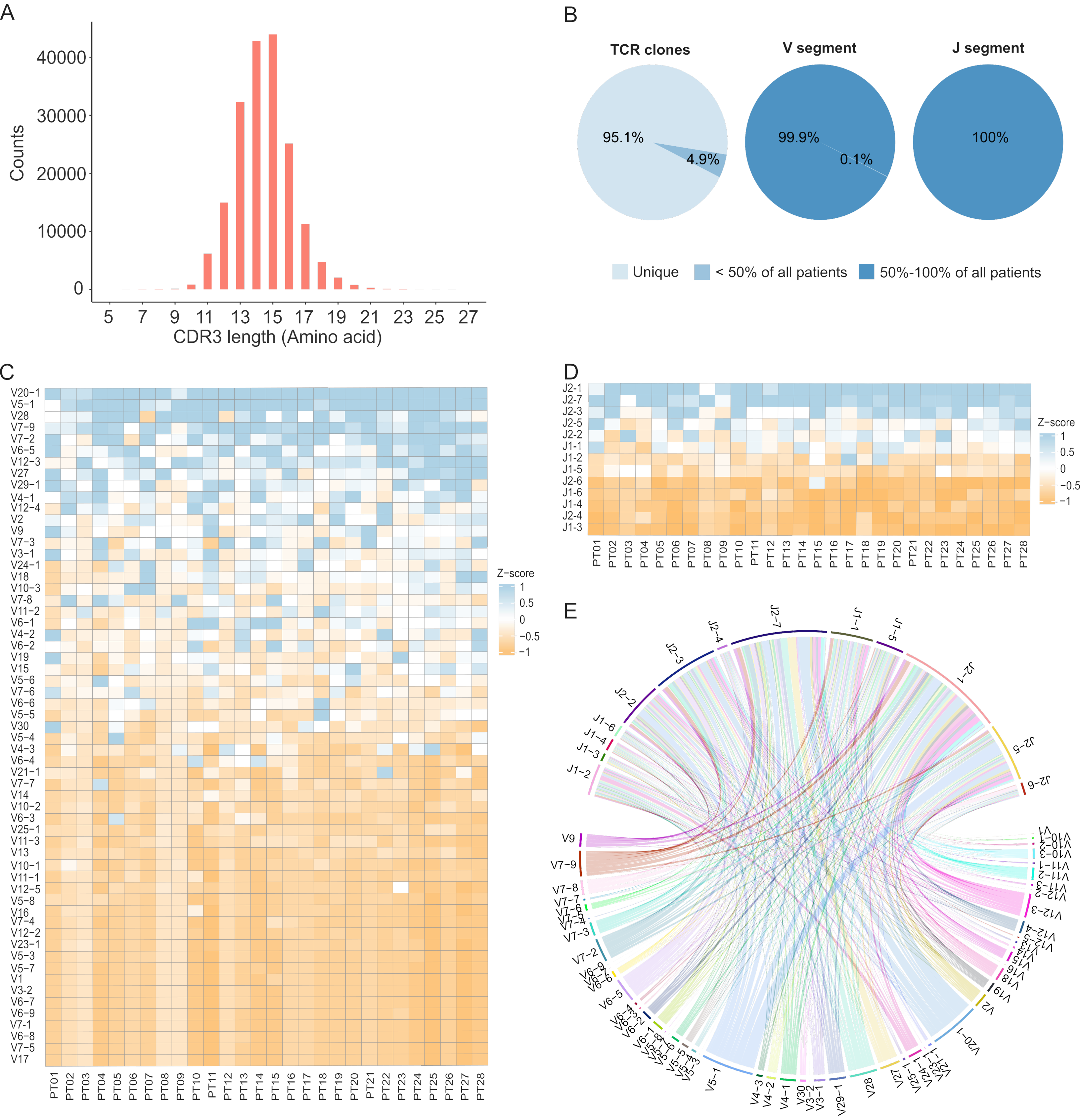

### Supplementary Figure 3

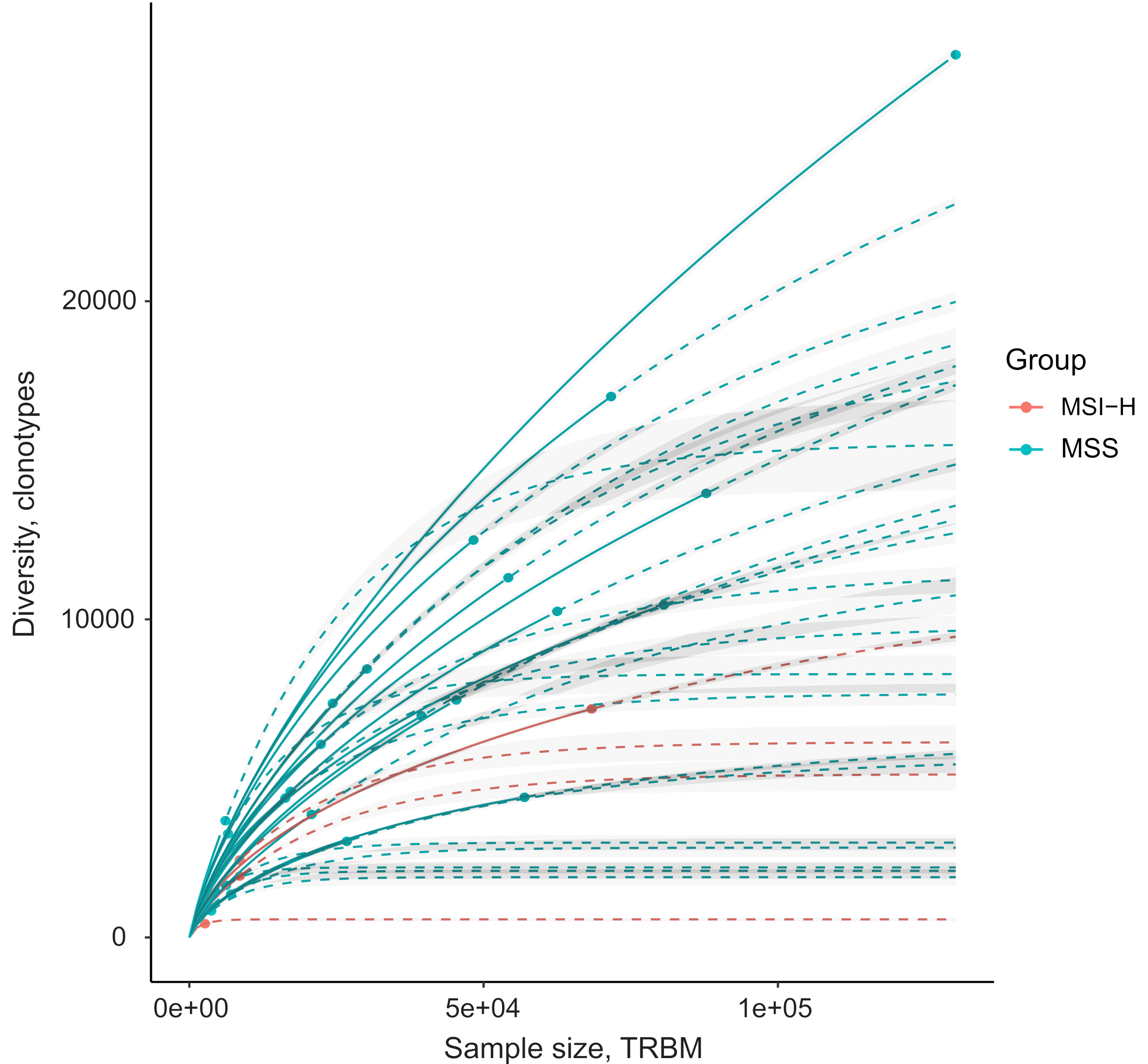

### Supplementary Figure 4

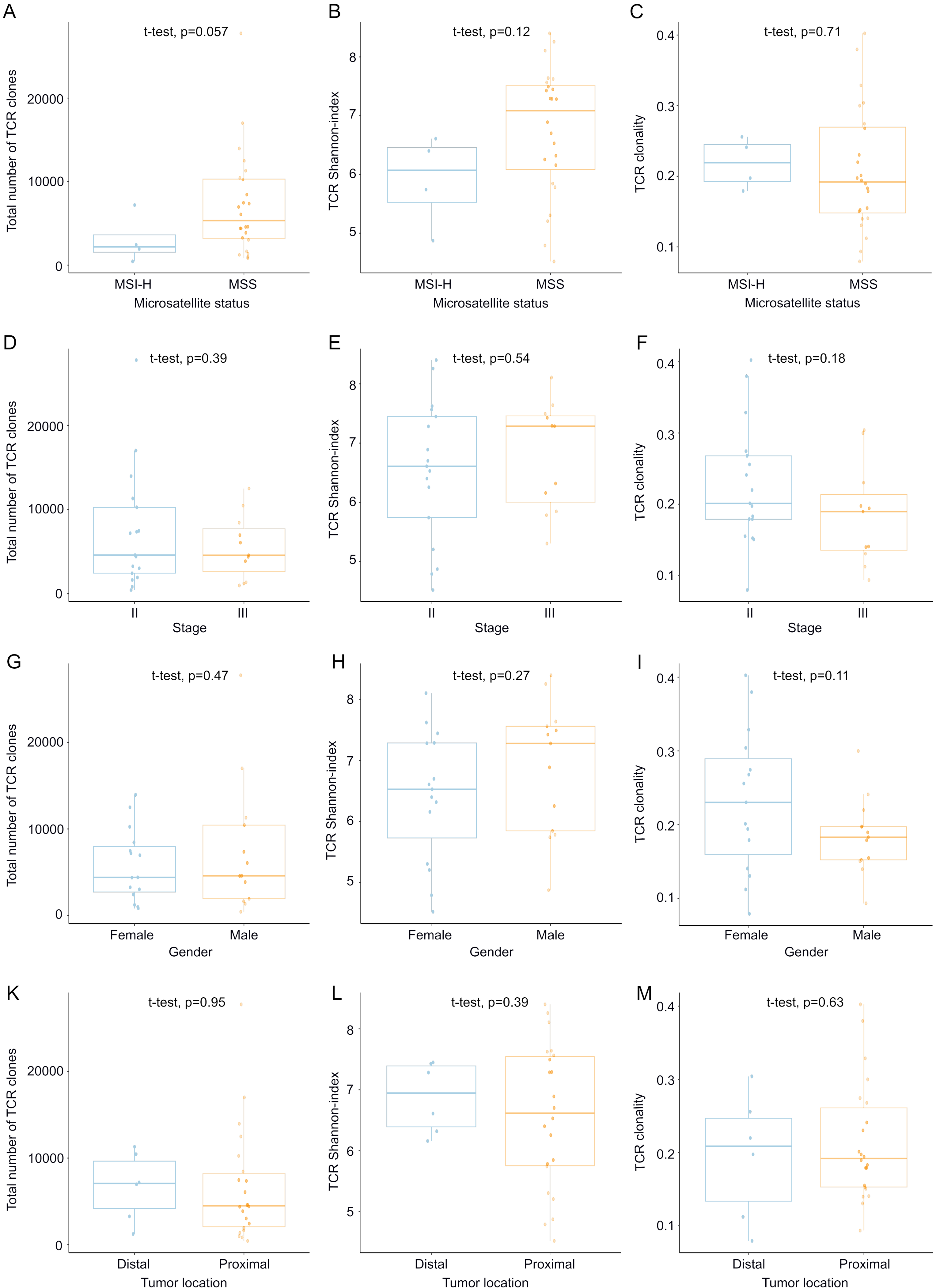

### Supplementary Figure 5

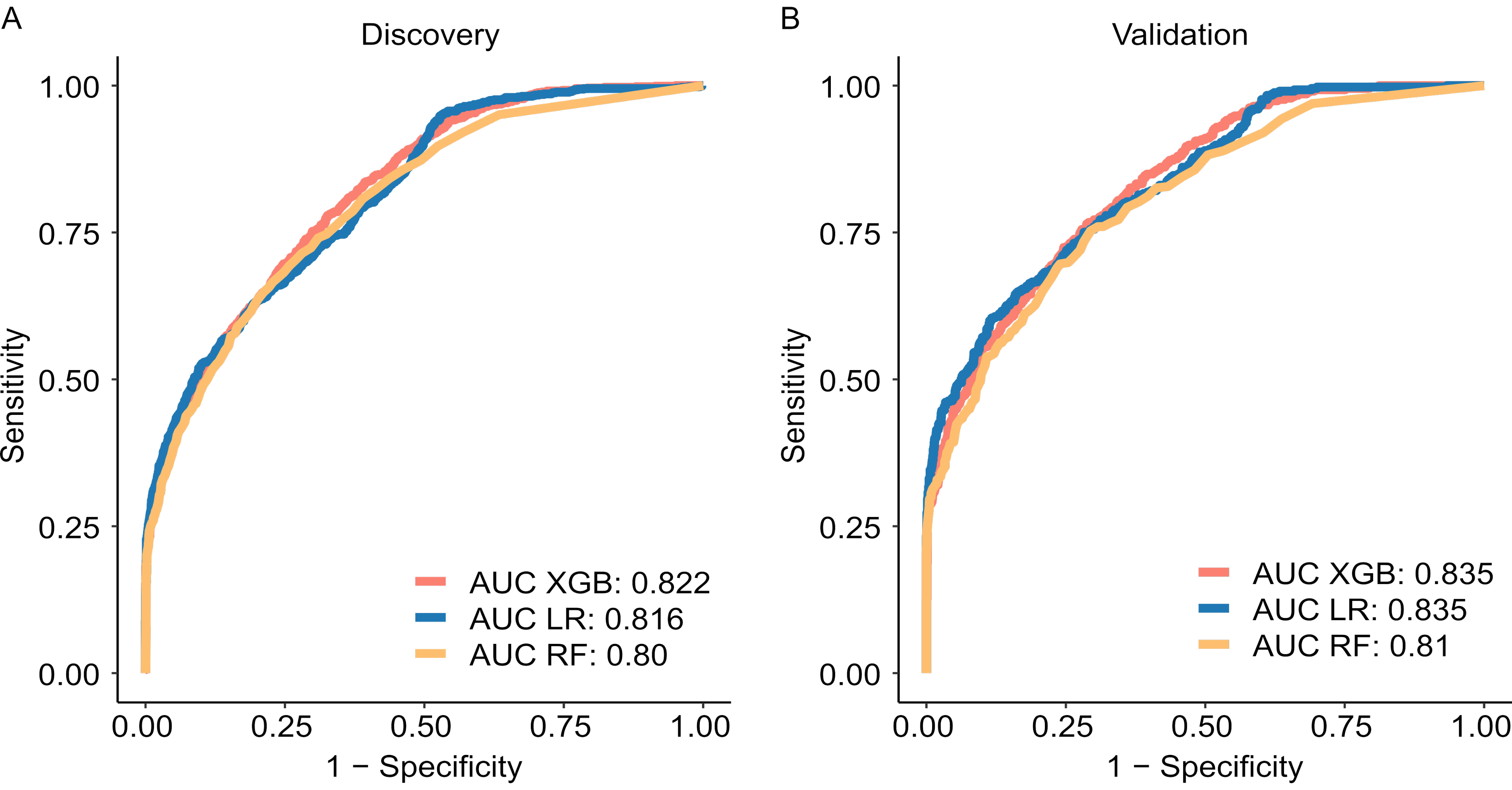

### Supplementary Figure 6

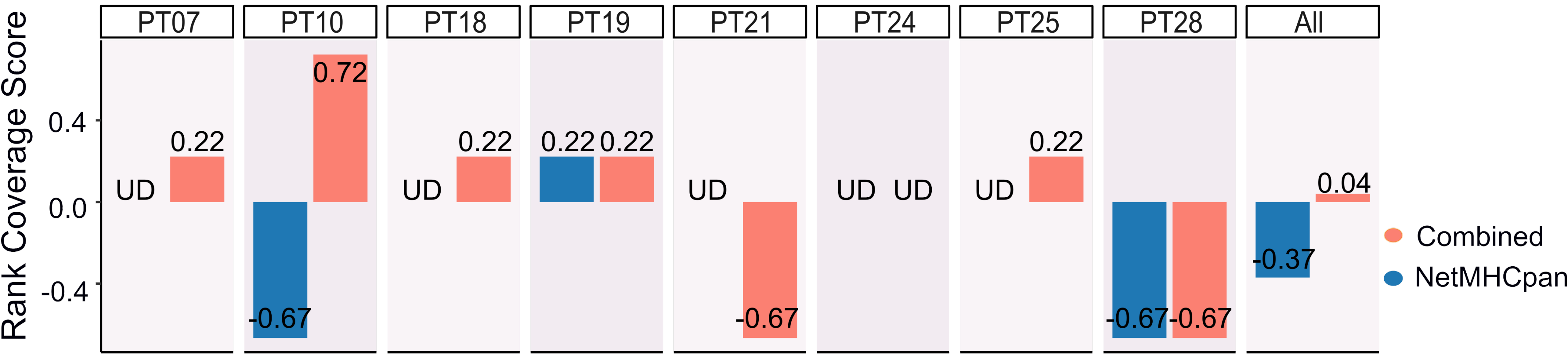
